# Dissociable oscillatory networks support gain and loss processing in human orbitofrontal cortex

**DOI:** 10.1101/2021.02.25.432874

**Authors:** Ignacio Saez, Jack Lin, Edward Chang, Josef Parvizi, Robert T. Knight, Ming Hsu

## Abstract

Human neuroimaging and animal studies have linked neural activity in orbitofrontal cortex (OFC) to valuation of positive and negative outcomes. Additional evidence shows that neural oscillations, representing the coordinated activity of neuronal ensembles, support information processing in both animal and human prefrontal regions. However, the role of OFC neural oscillations in reward-processing in humans remains unknown, partly due to the difficulty of recording oscillatory neural activity from deep brain regions. Here, we examined the role of OFC neural oscillations (<30Hz) in reward processing by combining intracranial OFC recordings with a gambling task in which patients made economic decisions under uncertainty. Our results show that power in different oscillatory bands are associated with distinct components of reward evaluation. Specifically, we observed a double dissociation, with a selective theta band oscillation increase in response to monetary gains and a beta band increase in response to losses. These effects were interleaved across OFC in overlapping networks and were accompanied by increases in oscillatory coherence between OFC electrode sites in theta and beta band during gain and loss processing, respectively. These results provide evidence that gain and loss processing in human OFC are supported by distinct low-frequency oscillations in networks, and provide evidence that participating neuronal ensembles are organized functionally through oscillatory coherence, rather than local anatomical segregation.

## Introduction

The human orbitofrontal cortex (OFC) is a critical node in reward-based decision-making: activity in OFC reflects value computations [1,2] and damage to OFC results in abnormal choice behavior [3,4]. Among the proposed functions of OFC, valuation and outcome processing are central. Valuation of positive and negative outcomes, which are necessary to learn about states of the world to inform future approach and avoidance behavior, have been associated with neural activity in OFC in both fMRI and animal studies [5–8].

Despite considerable progress, important questions remain regarding the organization of neuronal ensembles in valuation processes in OFC. In particular, while there is an increasing appreciation of the importance of neural oscillations in cognitive processing, whether they play a role in reward processing in OFC is unclear. Neural oscillations are generated by concurrent excitability fluctuations in groups of neurons, which generate periodic activity changes organized in several oscillatory bands (e.g. theta, 4-8 Hz, alpha 8-12 Hz and beta, 12-30Hz). Ongoing oscillations modulate input selection by favoring information that arrives at particular times in an oscillatory cycle, and allow the coordination of ensembles of neurons that share relevant information by establishing transiently synchronized networks [9]. Oscillatory coherence, in which oscillations across brain regions show a consistent phase relationship, is proposed to facilitate cross-areal communication by favoring phase-dependent activation of neurons [10]. Neural oscillations have been implicated in a variety of cognitive processes. In human studies, they have been extensively examined in non-invasive EEG and MEG studies and more recently in intracranial research (electrocorticography; ECoG). These studies have associated low-frequency neuronal oscillations with a variety of cognitive processes, including working memory [11,12], attention [13,14], sensory processing [15,16], motor control [17,18] and goal direction [19,20]. Prefrontal low-frequency oscillations have been specifically implicated in working memory, attention and spatial navigation [11,13,21,22]. Regarding reward processing, distinct scalp EEG frequency bands in prefrontal cortex have been shown to be differentially sensitive to gain and loss outcomes in the beta and theta bands, respectively [23,24].

Despite the proposed important of oscillatory processes in prefrontal function and the central role of OFC in reward-related processes, the nature of the involvement of oscillatory neural processing and coherence processes in reward processing in the human OFC remains poorly understood. This is in part due to due to the difficulty of measuring oscillatory activity in deep brain regions such as OFC using non-invasive approaches. Here, we leverage a unique patient population, human epilepsy patients undergoing intracranial monitoring, to directly examine the role of oscillatory neural activity in the human OFC during reward processing. Here we focus on low frequency (<30Hz) activity in a previous gambling task [25] to assess the association between oscillatory activity in OFC and reward processing.

Our results show that functionally distinct networks respond to gain and loss processing within OFC. Specifically, we observed a clear functional dissociation between sites in OFC associated with processing gain- and loss information, consistent with past EEG findings of frontal engagement in reward processing. However, unlike previous EEG findings, we found that monetary gains were associated with an increase in theta power (4-8Hz) whereas losses were associated with an increase in beta power (12-30 Hz) power. In addition, these frequency-specific power modulations were accompanied by selective increases in coherence, supporting the notion that reward-relevant information is organized in parallel neural ensembles oscillating in different frequencies. Anatomically, coherent theta/beta oscillations after gains/losses were not restricted to sites encoding gains/losses, indicating that coherence is an OFC-wide phenomenon. Finally, gain and loss networks were interspersed throughout the orbital surface, and did not follow any simple anatomical distribution (e.g., clusters or gradients). These results demonstrate that anatomically distributed low frequency oscillations differentially encode reward-related information in the human OFC, with power modulation in the theta and beta bands encoding gains and losses. In addition, gain and loss processing networks are not clustered anatomically. Instead, selective increases in OFC-wide oscillatory coherence suggest that these separate ensembles may be organized functionally through coherence. The combination of analysis of neural oscillations with decision-making models provides a novel approach to understand the neural basis of decision-making within and across brain areas.

## Results

We recorded LFP activity from 210 electrodes (192 after quality control; see Methods) in 10 patients while they played a gambling task (see Fig. 1A and Fig. S1 for electrode locations). Briefly, participants played 200 trials in which the choose between a sure prize and a risky gamble (Fig. 1B). Patient choices were dependent on gamble win probability, expected utility and risk (all p<10^-15^, random effects logit analysis) and was similar to that of healthy subjects (Fig. 1C, grey line; all comparisons p>0.2). Local field potentials (LFP) were recorded from all ECoG electrodes and frequency-band decomposed using a wavelet approach. We focus here on activity in low frequency bands (<30Hz); results from high-frequency analyses (70-200 Hz) were reported previously [26].

**Fig. 1.**
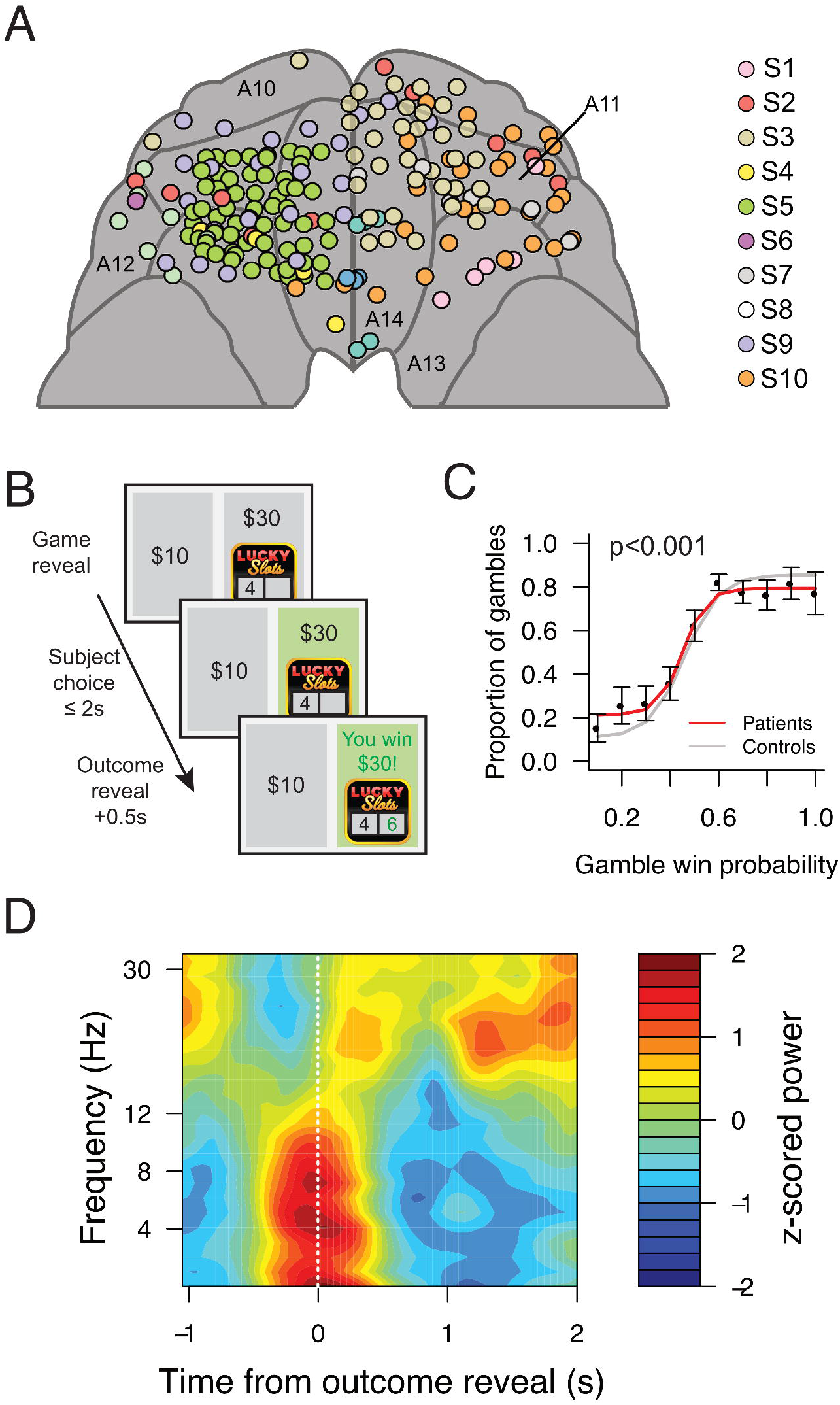
Experimental approach. *(A)* Anatomical reconstruction showing placement of ECoG electrodes (n=192) in OFC across all patients (n=10). Each color corresponds to a patient. Brodmann areas are indicated as A10/A11/A12/A13/A14. *(B)* Subjects (n=10) chose between a sure prize and a risky gamble with varying probabilities for potential higher winnings. Trials resulted in a win if a second number was higher than the first. Gamble outcome was shown regardless of choice. *(C)* Subjects’ choices were significantly affected by likelihood of winning the gamble (p<0.001, random effects logit analysis), and were comparable to choices of healthy controls (grey line; all p>0.2). (D) Power modulation associated with gamble outcome reveal across OFC sites. Plot indicates z-scored power modulation across frequencies (1-30Hz), relative to the patient choice to gamble or not (t=0). Gamble outcome reveal was at 550ms post-choice.

### Dissociation modulation of OFC low-frequency activity by outcome valence

To characterize whether power modulation in low frequency bands (theta, 4-8Hz; alpha, 8-12Hz; beta, 12-30Hz) carried relevant reward-related information during outcome evaluation, we generated a time-frequency representation (TFR) or neural activity using a wavelet approach. We observed power modulation across the theta and beta frequency bands, time locked to the reveal epoch (0-1.5 post-outcome reveal; Fig. 1D and S2). We then carried out linear regressions to identify gain/loss power modulation in each time-frequency tile. Specifically, we examined how much variance in neural power (percentage of explained variance, %EV) could be explained by gain/loss regressors across time and frequency bands (see Methods).

This generated Event-Related Computational Profiles (ERCPs) containing time x frequency depictions of the level of association between power and regressors of interest, which reveal the frequency specificity and timing of information encoding. We first examined the association between power encoding and gain events (i.e. trials in which the subject opted to gamble and won; Fig. 2) by averaging ERCPs across all patients and electrodes in our sample (n=192). We observed a significant association between gains and power in the delta-theta frequency bands (1-8Hz; Fig. 2A and C). Because slow oscillations in the delta band are difficult to estimate adequately given the duration of our analysis windows (~0.5-1s), we centered on analyzing the theta-band (4-8Hz). Fig. 2C shows an example electrode in which gain trials were associated with an increase in power compared to all other trials. Next, we performed a similar analysis for loss trials (Fig. 2B and D), which revealed a different activity pattern, with losses associated with modulation in the beta (12-30Hz) frequency band (Fig. 2B; individual electrode example in Fig. 2D). The average variance in the neural signal explained by gain outcomes was higher for the theta than for the beta frequency band, whereas the opposite was true for loss outcomes (Fig. 2E), indicating a double dissociation between beta-theta frequencies and outcome encoding. The direction of modulation was consistent across electrodes, with a majority showing increased theta power in gain trials (Fig. S4).

**Fig. 2.**
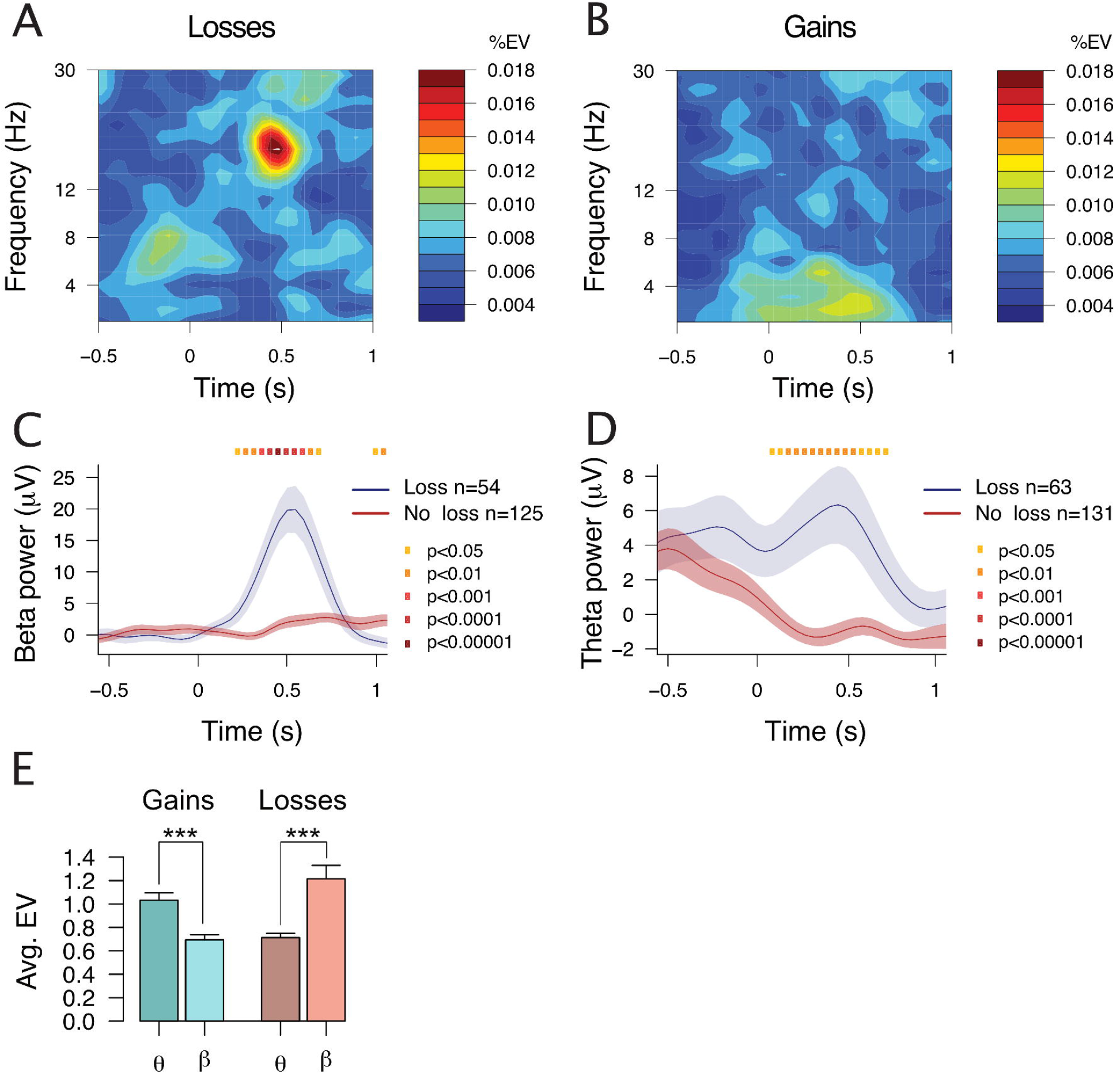
Distinct frequency band encoding of wins and losses. (A) and (B) Average event-related computational profile across all electrodes (n=192), indicating the strength of association (% explained variance, %EV) between loss (A)/win (B) outcomes and LFP power across frequency bands. Loss events are associated with beta power modulation, whereas win events are associated with delta/theta modulation. (C) Average beta power for loss/no loss trials from an example electrode encoding losses, separated by gamble outcome: loss (blue) or other outcomes (red). (D) as (C), but showing theta activity in a win encoding electrode for wins (blue) or other outcomes (red). (E) Average strength of association (% EV) between theta/beta band activity and gains (left) and losses (red).

To verify that these results were not driven by inter-subject or inter-electrode variation in neural activity, we used a nested mixed-effects model that included patient and electrode identity as random effects (see Methods). We found that regressions for both gains and losses were significantly active (p<10^-5^, corrected for multiple comparisons across frequency bands), indicating that gain/loss computations were robust across electrodes and patients. These results are consistent with a dissociable association between reward encoding in low frequency bands, with theta and beta band activity associated with gain and loss events, respectively.

### Overlapping anatomical distribution of gain and loss responses

Previous fMRI results have suggested an anatomical gradient of win/loss encoding, with loss responsivity higher in medial aspects of the OFC, and win processing located more laterally [7,27]. To examine whether there was anatomical segregation of gain- and loss-encoding in our ECoG dataset, we investigated the anatomical location of the encoding electrodes in our population.

We defined encoding electrodes as those showing a significant relationship between power modulation in beta (for losses) and theta (for gains) using a clustering approach followed by a permutation test (see Methods). Overall, we found that a similar proportion of electrodes encoded wins (30/192; 15.6%) and losses (29/192; 15.1%). A few electrodes encoded both losses and wins (6/192, 3.1%), but this proportion is not significantly overrepresented compared to a random overlap of both networks (p=0.58, chi-square test). Next, we examined the anatomical localization of these electrodes. To enable comparison across patients, patient scans and their corresponding electrode locations were normalized to template space (see Methods). We found that gain- and loss-encoding sets of electrodes were not segregated in distinct Brodmann areas, but instead were intermixed across the entire OFC surface (Fig. 3 and Fig. S4) suggesting anatomically distributed OFC encoding of gain and loss information.

**Fig. 3.**
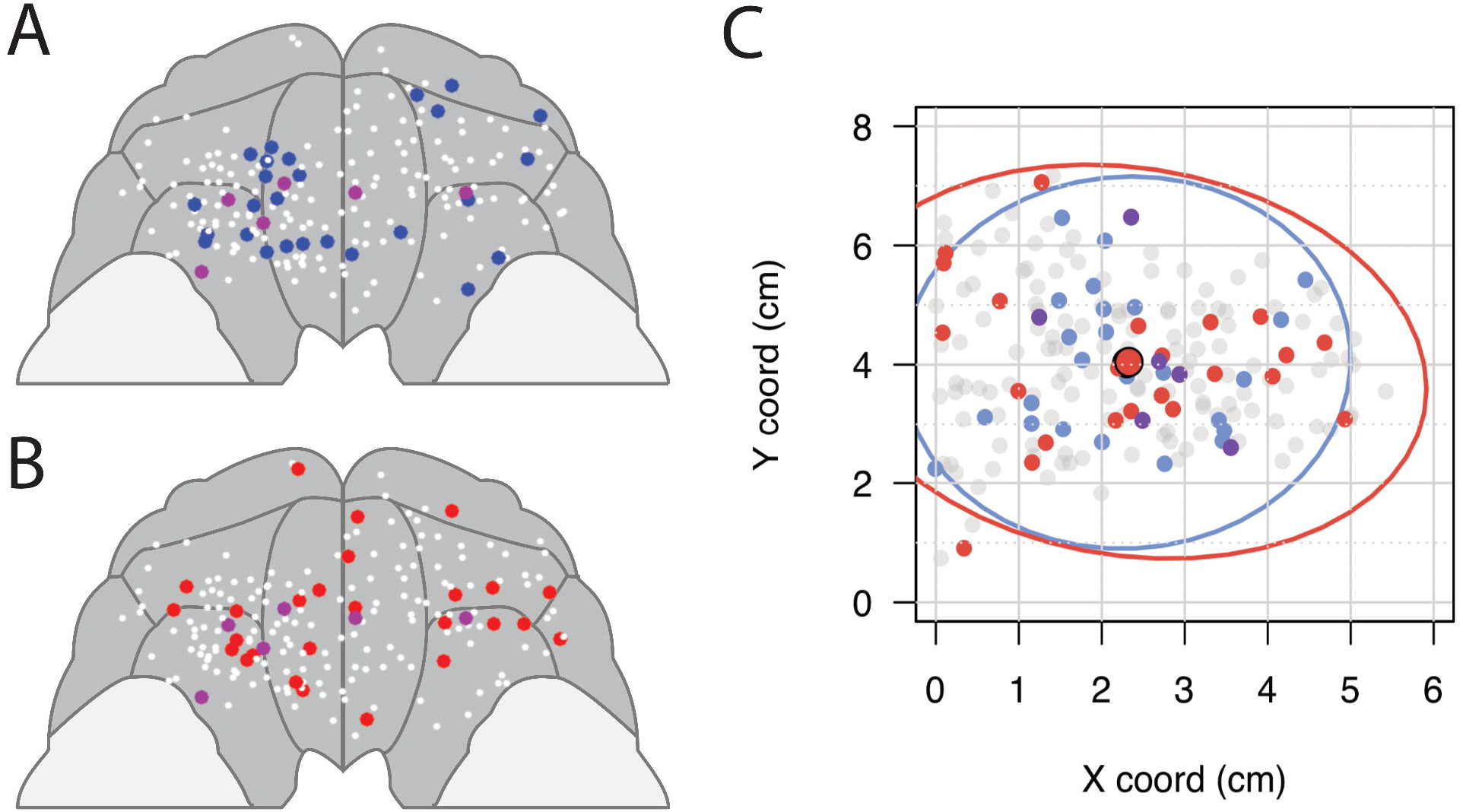
Overlapping gain and loss processing networks. (A) and (B): electrode positions projected on the orbital surface of a template brain. Electrodes are color-coded according to their reward-encoding characteristics: beta-band modulation in loss events (blue), theta-band modulation in win events (red), both (magenta) or no encoding (white). (C) Anatomical pattern of win and loss encoding. Scatterplot: X (medio-lateral) and Y (fronto-posterior) coordinates of all recorded electrodes, defined as distance from the z-projection of the anterior commissure (AC), on a single hemisphere. Color coding as in (A-B). Ellipses indicate 95% confidence interval across X and Y coordinates; centroids for loss and wins ellipses, indicated by the black-outlines, are overlapping.

### Neural activity in OFC shows outcome and frequency-specific coherence

Since cortical sites engaged in gain and loss processing are not anatomically clustered, another possibility is that they are instead organized as a functional ensemble through coordinated changes in inter-electrode coherence. Coherent neural oscillations have been proposed as a potential mechanism to achieve functional communication across cortical sites, which could play a role in sharing outcome-specific information across OFC sites. To assess whether neural oscillations were engaged during outcome processing, we examined low frequency coherence after outcome reveal. To assess this possibility, we calculated coherence at the time of the outcome reveal event across all low-frequency bands (1-30Hz) for all pairs of OFC electrodes for each patient in our dataset. To compensate for potential differences in baseline coherence across patients and electrodes, we used a within-electrode analytical strategy, calculating coherence separately for different trial types (loss, win and safe bet) after the gamble outcome reveal. The resulting coherence estimates were then compared using a mixed-model approach (see Methods) that include electrode and patient identity as random effects terms.

The results showed that win and loss events were accompanied by significant increases in coherence, in a frequency-specific manner consistent with the power encoding results. Specifically, gain events were accompanied by an increase in theta-band coherence (Fig. 4C), whereas loss events were associated with an increase in beta coherence (Fig. 4D). To verify that these coherence increases were not solely driven by an increase in power modulation, we conducted a linear regression analysis in which we examined the association between the average power across electrodes in each pair and their coherence, separately for each frequency band (theta/beta). We found no evidence that higher coherence was associated with higher power values across electrode pairs (both p>0.3), revealing that the coherence effects were separable from the power modulation effects. As was the case with the power results, there was a double dissociation between gain/loss outcome encoding and frequency-specific coherence increases (Fig. 4E). Finally, direct comparison of the time-courses of gain-theta and loss-beta power and coherence modulation showed comparable time profiles and onsets (Fig. S5).

**Fig. 4.**
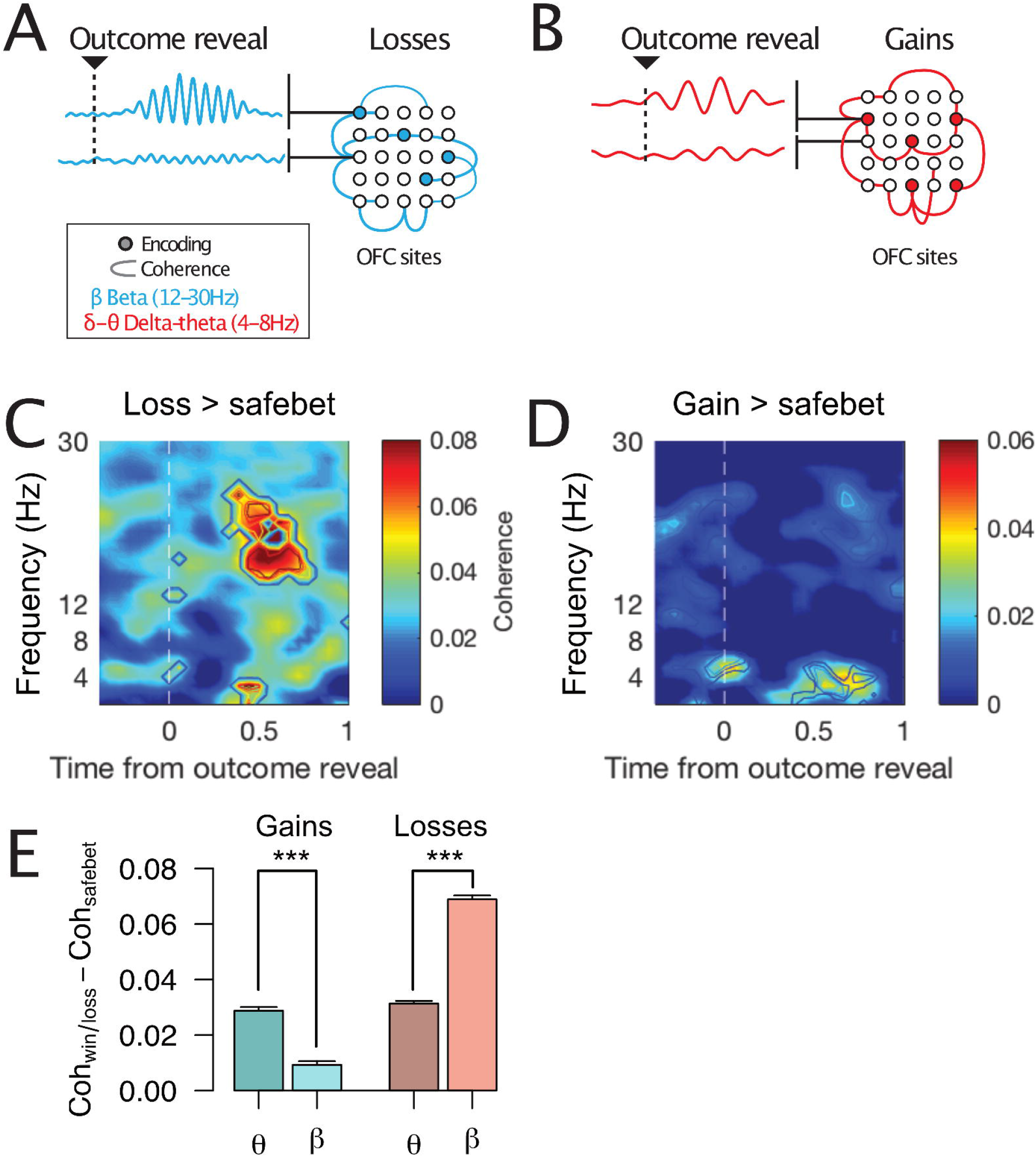
Oscillatory coherence organizes network of active/inactive cortical sites. (A) Cartoon depicting the power/coherence modulation results. Losses are associated with beta (12-30Hz) power increases in a number of cortical sites (blue dots), which engage in beta coherence with other encoding/non-encoding sites (blue lines). (B) As (A), but for gain encoding. The results are quantitatively similar to gain encoding, but the set of encoding cortical sites is different, and power/coherence modulation is in the theta (4-8Hz) frequency band. (C) Average difference in coherence between loss and safebet trials across all pairs of electrodes. The white vertical dotted line at t=0 indicates gamble outcome reveal. Contour lines indicate statistical significance (p<0.05, p<0.01, p<0.001, etc.) as established by a mixed-model analysis. (D) As (C), but for gain vs safebet trials. (E) Overall differences in coherence for gains (left) and losses (right) in the theta and beta frequency bands, showing a dissociable theta-gains and beta-losses association.

### Comparison to HFA results

In a previous study, we described encoding of reward-related information in high-frequency activity (HFA) in human OFC [26]. Because gain and loss information was also reflected in HFA, we examined the relationship between HFA and low-frequency encoding in human OFC by directly comparing both sets of results. First, we compared the proportion of cortical sites encoding gains and losses in low and high-frequencies. We found that a comparable number of cortical sites encoded theta-gain (n=29/192 electrodes) and beta-loss (n=30/192 electrodes), and comparable to the proportion of HFA-gain (n=45/192) and HFA-loss (n=33/192) encoding sites we reported earlier. Thus, outcome encoding in beta/theta and HFA recruited activation in a comparable number of cortical sites.

One possibility is that low and HFA encoding reflect activation of the same network of cortical sites. To examine whether that was the case, we next examined the proportion of cortical sites encoding outcomes in both HFA and low frequencies. We found that electrodes encoding gains in both HFA and theta band activity were not overrepresented (n=9/192 vs 8.15/192 expected from random mixing, p=0.13, χ^2^ test). However, electrodes encoding in both HFA and beta were slightly overrepresented (n=9/192 observed vs 4.53/192 expected from random mixing, p<0.05, χ^2^ test). Overall, these results do not provide strong support for the notion that modulation in both low frequencies and HFA occurs in the same cortical sites.

## Discussion

We assessed whether neuronal oscillations, implicated in a variety of cognitive processes in human prefrontal cortex, play a role in processing reward outcomes in the OFC. To test this notion, we carried out multi-electrode ECoG recordings directly from the OFC of human neurosurgical patients while they made a series of decisions under uncertainty in gambling game. Our results show that low-frequency neural activity in human OFC encodes information about reward outcomes, with losses and gains having separable physiological and anatomical substrates. Specifically, we found that gains were associated with power increases in theta-band (4-8Hz), whereas losses were associated with power increase in the beta band (12-30Hz). Cortical sites showing significant theta/beta power modulation were not anatomically segregated, but rather interspersed across the orbitofrontal surface. Finally, we observed a concomitant increase in OFC-wide coherence in theta (for gains) and beta (for losses) that was not driven by power increases.

### Oscillations encode reward-related information

Prefrontal low-frequency oscillations have been implicated in a variety of cognitive processes, including working memory, attention, language and spatial navigation [11,13,21,22,28]. Here, we add to the growing body of work implicating neural oscillations in reward outcome processing in decision-making [23,24]. EEG studies also suggest a differential role for low-frequency bands in reward processing, with theta and beta-band activity as the main oscillatory substrates for gain and loss processing, but the nature of their association varies across studies. For instance, increases in beta power were associated with gain processing [23,29,30], and increases in theta power with losses or negative feedback [23,31]. MEG recordings in humans have also proposed an increase in theta OFC is associated with win outcomes [32]. However, other EEG studies shows a reverse pattern more consistent with the one we report here, with beta activation in response to no reward or error conditions (comparable to our loss trials) and theta in reward conditions [24,33].

There are several possible explanations for these discrepancies. One possibility is that they are due to methodological differences between EEG/MEG/ECoG. However, even within modality, opposite effects can be found (e.g. EEG [30,33]), which makes this unlikely to be the only source of discrepancy. Another possibility is that there are differences in the source of oscillatory activity. EEG sources vary between prefrontal (Fz) and lateral (F6), with common estimated anatomical locations in ACC and LPFC, but they are unlikely to capture activity originating from the OFC. However, given that these areas are all implicated in reward processing [5,8,34,35] and the proposed role of oscillations in establishing functional connectivity across brain areas [10], it is possible that LPFC/ACC and OFC oscillations are functionally related. If that was the case, discrepancies between OFC and LPFC/ACC oscillations may reflect the different roles (bottom-up vs top-down) established by both reward-responsive areas, or their relative temporal organization. For example, reward information may be processed first in OFC and then communicated to LPFC, and this functional communication may be reflected in oscillatory activity or in temporal activation patterns.

Another possible explanation relates to the different definitions of gains and losses across tasks. In some EEG studies, losses refer to a negative money gain [23] (but see [24]). In our study, loss events can also be described as absence of reward, rather than an actual loss (e.g. negative gains), a design that we adopted because of limitations, including human subjects protections and working with patients. Thus, encoding of losses may be associated with a different neural representation altogether. Finally, it is also possible that they reflect different cognitive demands across tasks. Unlike some of the non-invasive experiments, our gambling task does not require working memory. Since working memory load is associated with theta band power [36], it is possible that the different cognitive demands of a working memory-reward task, as compared to a gambling task with no memory demands, engages another circuitry indexed by different oscillatory mechanisms. Despite these discrepancies, these different studies consistently show that neural activity across theta and beta bands is recruited during feedback processing, and that they represent separate neural processing channels for gains and losses.

### Functional organization of low frequency responses to gains and losses

Neuroimaging data suggests anatomically-segregated processing for different aspects of reward in OFC, for options varying in desirability (appetitive/aversive), abstractness (primary rewards such as food, water vs. secondary rewards such as money), as well as for valuation/choice processes [7,27,37,38], suggesting the existence of anatomical segregation of reward functions. Our data, however, does not support anatomical segregation of gain and loss low frequency encoding across OFC subregions in our task (Fig. S3). Rather, we observed that win and loss networks were distributed across the orbitofrontal surface (Fig. 4), which is also consistent with a similar pattern with reward-related HFA encoding [26], suggesting that separation of reward information is not associated with anatomical segregation.

Instead, coordination of distinct types of reward across cortical sites may be supported by functional activity patterns manifesting as oscillatory coherence. In support of this hypothesis, we observed a generalized increase in low-frequency coherence across OFC electrodes during reward outcome. Consistent with the power modulation results, this coherence increase was frequency- and outcome-specific, with separate beta and theta coherence increases following losses and gains, respectively (Fig. 4).

There are several potential functional roles for these coherence increases. Low frequency oscillations in monkey PFC reflect abstract rules relevant to ongoing behavior, with beta and alpha synchronies organizing neural ensembles representing different rules [20]. A similar mechanism may be at play here, with theta and beta ensembles carrying complementary but distinct reward information. In addition, coherence has been proposed to support functional communication across cortical sites [10]. Thus, one possibility is that synchronous oscillations in OFC facilitate information sharing across cortical sites with distinct encoding properties. Furthermore, we observed that coherence increases are not driven by power increases. For example, loss (gain) events, which are associated with an increase in beta (theta) power, may entrain other OFC sites through oscillatory coherence without directly modulating their oscillatory power. Coherent oscillations may also provide a mechanism to broadcast reward relevant information from OFC to other reward-responsive cortical areas (e.g. LPFC), reflecting cross-areal information processing, an idea that will need to be tested in future experiments.

Overall, the existence of these separate oscillatory networks, in addition to the diffuse anatomical organization described above, supports the notion that parallel neural ensembles carry different but complementary types of reward-related information. Interestingly, the increase in theta coherence was not limited to the time of outcome reveal, but showed a peak that ramped up before gamble reveal (Fig. S3). Given that patients choose to gamble more often in trials in which win probability is higher (Fig. 1C), expectation of reward is correlated with win outcomes in our dataset. Thus, it is possible that this pre-reveal activation reflects a reward expectation effect. Consistent with this idea, neurons in the orbitofrontal cortex of rodents have been shown to phase-lock to theta band oscillations in anticipation of reward [39], and theta-band activity is also modulated in human frontal cortex [40]. Theta is also related to attentional processes in other cortical regions, compared to a role of beta in top-down processing [41], so an alternative explanation would be an increase in attention in trials in which a positive outcome is expected.

### Separate oscillatory mechanisms for gain and loss processing – functional relevance

The existence of separate, but related encoding mechanisms across different frequency bands for gain and loss processing in the human OFC raises a number of questions on their neurobiological origin and functional significance. Low frequency LFP activity captures a diverse number of voltage generators, the most prominent ones being postsynaptic currents, both excitatory and inhibitory. In contrast, activity in higher frequency bands (60Hz and above, i.e. high frequency activity; HFA) reflects local cortical activation, including neuronal spiking and dendritic currents [42,43]. Thus, it is possible that encoding in low frequency and HFA may capture different activation aspects of the same neuronal ensembles. For example, theta/beta activation may reflect input to OFC cortical sites, with HFA reflecting spiking output of the same neuronal population. If these processes reflected different aspects of activation of a single neuronal population, we would expect significant overlap between low-frequency and HFA encoding sites. However, in our dataset the amount of overlap was modest, with only a slight overrepresentation of concurrent HFA and beta-loss encoding, and none for theta-gain encoding. These observations suggest that low-frequency and HFA encoding mechanisms do not simply reflect different activation aspects of the same neuronal population. One possibility is that localized low-frequency input modulates activity throughout other OFC sites by modulating the degree of oscillatory coherence (Fig. 4), which could facilitate synchronous spiking in entrained sites and information propagation to downstream targets [44].

The question arises on the specific roles of beta and theta frequency bands in reward processing. One possibility is that they reflect different information processing streams or cognitive processes. Different oscillations may index the engagement of distinct downstream targets, reflecting the need for different adaptive behavioral strategies after gain/loss events. If this was the case, one can imagine loss events favoring a strategy change after adverse events (i.e. ‘switch’), with win events favoring perseverance and continued attention after reward (i.e. ‘stay’). However, previous evidence suggests this may not be the case: beta-band activation has been proposed to play a role in maintaining, rather than altering, ongoing behavioral patterns [41]. Consistent with this, electrical stimulation of the caudate nucleus results in extraneous modulation of beta-band activity and repetitive OCD-like behavior and negative affective states in macaques [45]. Alternatively, the engaged theta/beta networks could be related not to behavioral, but to emotional responses after positive/negative events. Consistent with this idea, prefrontal human beta rhythms, including the ventromedial prefrontal [46] and anterior cingulate cortices [47] have been implicated in emotional processing and mood regulation. In addition, beta coherence in limbic areas (hippocampus and amygdala) has been associated with mood in human patients [48], and different frequency bands in the amygdala-hippocampal circuit underlie separation of emotionally relevant information [49]. In the context of our decision-making task, unexpected losses are expected to have a negative emotional impact [50]. Thus, prefrontal beta oscillations may be a general mechanism underlying emotional responses to negative outcomes in prefrontal and limbic regions.

### Conclusion

Here we demonstrate that neural oscillations in the human OFC encode behaviorally relevant reward information, with anatomically interspersed and functionally distinct networks in OFC encoding positive (gains) and negative (losses) outcomes indexed by power modulations in the theta and beta bands, respectively. These network-specific power modulations were accompanied by OFC-wide oscillatory coherence in the theta band and reward and the beta band in loss, providing a potential mechanism for establishment of rapid and reversible functional connectivity at behaviorally relevant time points. Thus, reward engages separate OFC rhythms associated with the establishment of distinct brain networks for adaptive decision-making behavior.

## Materials and Methods

### Subjects

Data was collected from 10 (4 female) adult subjects with intractable epilepsy who were implanted with chronic subdural grid and/or strip electrodes as part of a pre-operative procedure to localize the epileptogenic focus. We paid careful attention to the patient’s neurological condition and only tested when the patient was fully alert and cooperative. The surgeons determined electrode placement and treatment based solely on the clinical needs of each patient. Data were recorded at four hospitals: the University of California, San Francisco (UCSF) Hospital (n=2), the Stanford School of Medicine (n=2), the University of California, Irvine Medical Center (UCI) (n=5) and at Albany Medical College (n=1). Due to IRB limitations, subjects were not paid for their participation in the study but were encouraged to make as many points as possible. As part of the clinical observation procedure, patients were off anti-epileptic medication during these experiments. Healthy participants (n=10) with no prior history of neurological disease were recruited from UC Berkeley’s undergraduate population and played an identical version of the gambling task. All subjects gave written informed consent to participate in the study in accordance with the University of California, Berkeley Institutional Review Board.

### Behavioral task

We probed risk-reward tradeoffs using a simple gambling task in which subjects chose between a sure payoff and a gamble for potential higher winnings. Trials started with a fixation cross (t=0), followed by the game presentation screen (t=750ms). At that time, patients were given up to 2s to choose between a fixed prize (safe bet, $10) and a higher payoff gamble (e.g. $30; Figure 1). Gamble prizes varied between $10 and $30, in $5 increments. If the patient did not choose within the allotted time limit, a timeout occurred and no reward was awarded for that round. Timeouts were infrequent (9.98% of all trials) and were excluded from analysis. Gamble win probability varied round by round; at the time of game presentation, subjects are shown a number between 0-10. At the time of outcome (t=550ms post-choice), a second number (also 0-10) is revealed, and the subject wins the prize if the second number is greater than the first one. Only integers were presented, and ties were not allowed; therefore, a shown ‘2’ had a win probability of 80%. The delay between buttonpress and gamble outcome presentation (550ms) was fixed, and activity for both epochs is temporally aligned. Therefore, offer value, risk and chosen value vary parametrically on a round-by-round basis, and patients had full knowledge of the (fair) task structure from the beginning of the game. Both numbers were randomly generated using a uniform distribution. The gamble outcome (win/loss) was revealed regardless of subject choice, allowing us to calculate experiential and counterfactual prediction errors (see Behavioral analysis, below). A new round started 1s after outcome reveal. Patients played a total of 200 rounds (plus practice rounds), and a full experimental run typically lasted 12-15min. Location of safe bet and gamble options (left/right) was randomized across trials. Patients completed a training session prior to the game in which they played at least 10 rounds under the experimenter’s supervision until they felt confident they understood the task, at which point they started the game. This gambling task minimized other cognitive demands (working memory, learning, etc.) on our participants.

### ECoG Recording

ECoG was recorded and stored with behavioral data. Data collection was carried out using Tucker-Davis Technologies (Albany, Stanford and UCSF) or Nihon-Kohden (at UCI) systems. Data processing was identical across all sites: channels were amplified x10000, analog filtered (0.01-1000 Hz) with >2kHz digitization rate, re-referenced to a common average off-line, high-pass filtered at 1.0 Hz with a symmetrical (phase true) finite impulse response (FIR) filter (~35 dB/octave roll-off). Channels with low signal-to-noise ratio (SNR) were identified and deleted (i.e. 60 Hz line interference, electromagnetic equipment noise, amplifier saturation, poor contact with cortical surface). Out of 210 OFC electrodes, 192 were artifact-free and included in subsequent analyses. Additionally, all channels were visually inspected by a neurologist to exclude epochs of aberrant or noisy activity (typically <1% of datapoints). A photodiode recorded screen updates in the behavioral task, recorded in the electrophysiological system as an analog input and used to synchronize behavioral and electrophysiological data. Data analysis was carried out in MATLAB and R using custom scripts.

### Electrophysiological analysis

ECoG recordings were downsampled to 1KHz. Channels were visually examined and those with low quality recordings due to bad electrode-brain contact were excluded from analysis. In our patient sample, no epileptic electrodes were located in OFC. Recordings were visually examined by a neurologist (RTK), and any trials containing aberrant epileptiform activity were excluded from subsequent analysis. Electrodes were then re-referenced using a within-grid/strip common average reference (CAR). Time-frequency decomposition was carried out using a multitaper approach. Briefly, whole-recording spectrograms were created for each electrode using log-spaced frequencies between 1 and 30Hz. Spectrograms were then subset by selecting windows of interest around outcome events, as indicated by the behavioral timestamps, and baseline-subtraction was carried out for each frequency of interest. For the trials in figure 1C and D, power was calculated by averaging for theta (4-8Hz) and beta (12-30Hz) across log-spaced frequency bins (4 and 11 frequency bins, respectively).

### Behavioral Analysis

We classified outcomes as win/loss/safe bets, depending on the patient choice and gamble outcome. Gains and losses refer to gamble trials; safe bet trials refer to trials in which the patient decided not to gamble, regardless of subsequent gamble outcome. To examine the relationship between power modulation and win/loss events, we used a linear regression approach. For each frequency and time of interest, we regressed the power estimate against outcome (win/loss). The resulting R^2^ was then presented as a time-frequency event-related computational profile (ERCP; figures 1A-B) representing the association between power modulation and the regressor of interest.

### Coherence analyses

Cross-electrode coherence was calculated using the Fieldtrip toolbox [51]. For each within-patient pairwise electrode combination, time-frequency decomposition was carried out using a Hanning window for frequencies between 1 and 30Hz. Coherence analysis was carried out using the <u>ft</u>_connectivityanalysis function, separately for loss, win and safe bet trials for each electrode pair and frequency band. To account for inter-subject variability, we compared the coherence values between loss (Fig. 4A) and win (Fig. 4B) events and safe bet events by using a mixed-effect model that includes subject and electrode identity as random effects. We used the mixed model to analyze the relationship between coherence and trial type (win/loss) for all frequency-time combinations in the time immediately preceding and subsequent to outcome reveal, and captured the statistical significance results as time-frequency contours.

Because of the limitations associated with coherence analyses (i.e. they must be carried out in a within-patient basis, and involve pairs of electrodes which limits its power in patients with lower number of electrodes), limiting coherence analyses to pairs of encoding electrodes would have resulted in a small number of pairs, and limited statistical power. Thus, we instead chose to examine coherence across all electrode pairs for each patient.

### Anatomical reconstructions

For each patient, we collected a pre-operative anatomical MRI (T1) image and a post-implantation CT scan. The CT scan allows identification of individual electrodes but offers poor anatomical resolution, making it difficult to determine their anatomical location. Therefore, the CT scan was realigned to the pre-operative MRI scan. Briefly, both the MRI and CT images were aligned to a common coordinate system and fused with each other using a rigid body transformation. Following CT-MR co-registration, we compensated for brain shift, an inward sinking and shrinking of brain tissue caused by the implantation surgery. A hull of the patient brain was generated using the Freesurfer analysis suite, and each grid and strip was realigned independently onto the hull of the patient’s brain. This step often avoided localization errors of several millimeters. Subsequently, each patient’s brain and the corresponding electrode locations were normalized to a template using a volume-based normalization technique, and snapped to the cortical surface [52]. Finally, the electrode coordinates are cross-referenced with labeled anatomical atlases (JuBrain and AAL atlases) to obtain the gross anatomical location of the electrodes, verified by visual confirmation of electrode location based on surgical notes. Only electrodes confirmed to be in OFC (n=192) were included in the analysis. For display purposes, electrodes are displayed over a traced reconstruction of the ventral surface showing putative Brodmann areas. For analysis of anatomical location of encoding electrodes (Fig. 3), we defined beta-loss and theta-gain encoding electrodes as those that showed a significant association as indicated by a permutation test. Briefly, to leverage the time profile of the signals without imposing restrictions on activation timing, an aggregate statistic was calculated as the sum of F-stats for the longest stretch of consecutive windows showing a significant association between power and win or loss (linear regression p<0.05). The aggregate F-stat was subject to a permutation test by shuffling the behavioral labels (n=1,000 permutations). We then took the proportion of permuted fits with a sum-of-F-stat higher than that in the original dataset as the permutation p-value, which was further corrected using a Bonferroni correction (across n=192 electrodes). Electrodes with a corrected permutation p-value <0.05 were considered active.

## Acknowledgements

We would like to thank the patient volunteers and the research and surgical staff at the recording sites for their support and cooperation, and members of the Knight lab for assistance with data collection. This project was supported by NINDS R37NS21135, DARPA SUBNETS UC (to RTK), NIMH MH112775 (to MH), NIMH K01MH108815 (to IS) and R21MH109851 (to RTK, MH and IS).

## Notes

### Competing Interest Statement

The authors have declared no competing interest.

