## Supplementary online materials for "Dissociable oscillatory networks support gain and loss processing in human orbitofrontal cortex"

**Supplementary online material**

Fig. S1: Anatomical coverage. Individual reconstruction of the 10 patients included in the study, showing location of all ECoG electrodes in the ventral surface of the brain. The final number of electrodes, after excluding those located outside of OFC and with aberrant activity, was 192.

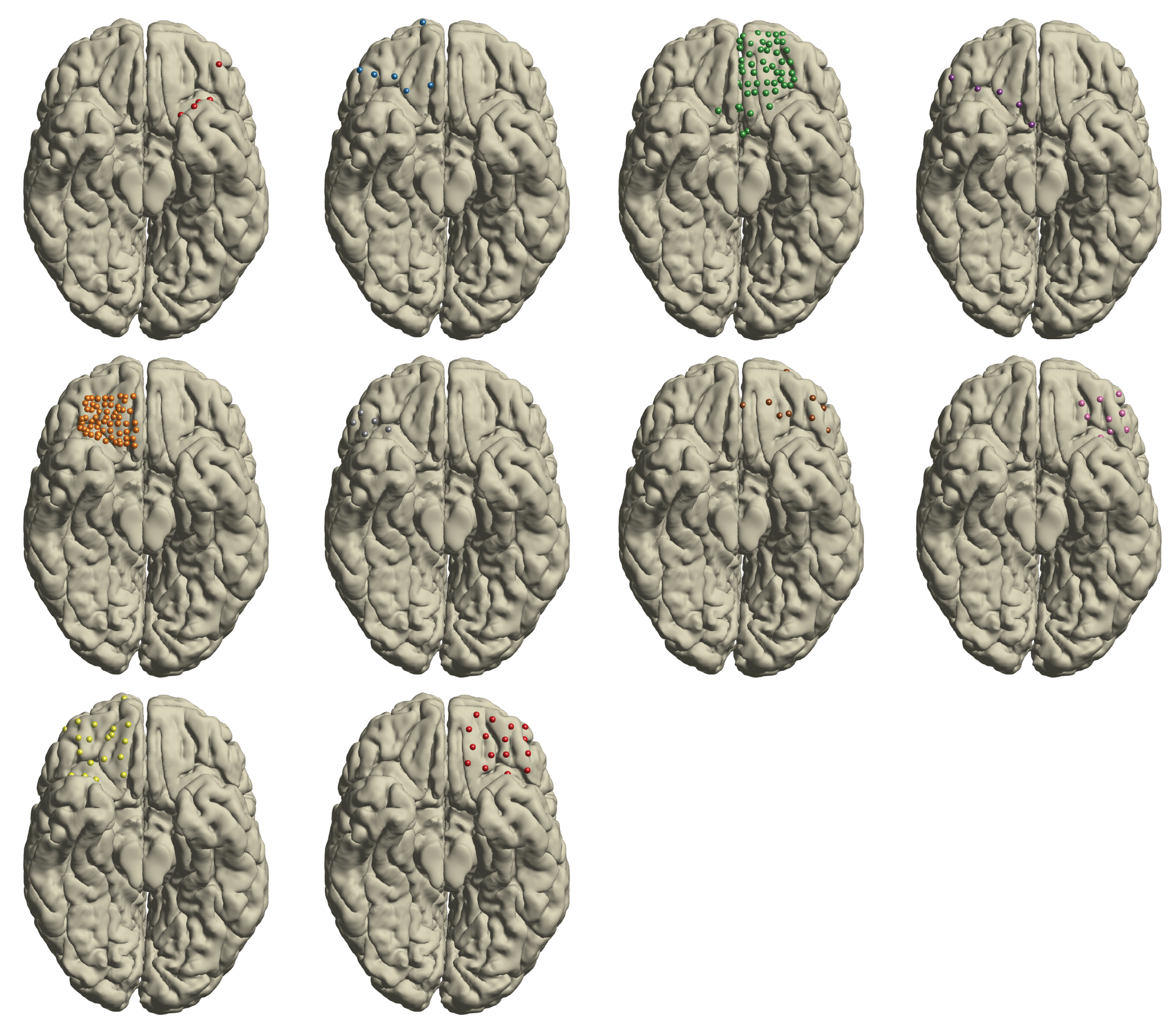

**Fig. S2 – Single trial traces for electrodes encoding gains and losses.** (A) Beta power trials from an example loss encoding electrode, separated by gamble outcome: losses (top) or other outcomes (middle). (B) As (A), but for gain encoding in the theta band. Trials correspond to the same example electrodes in Fig. 1

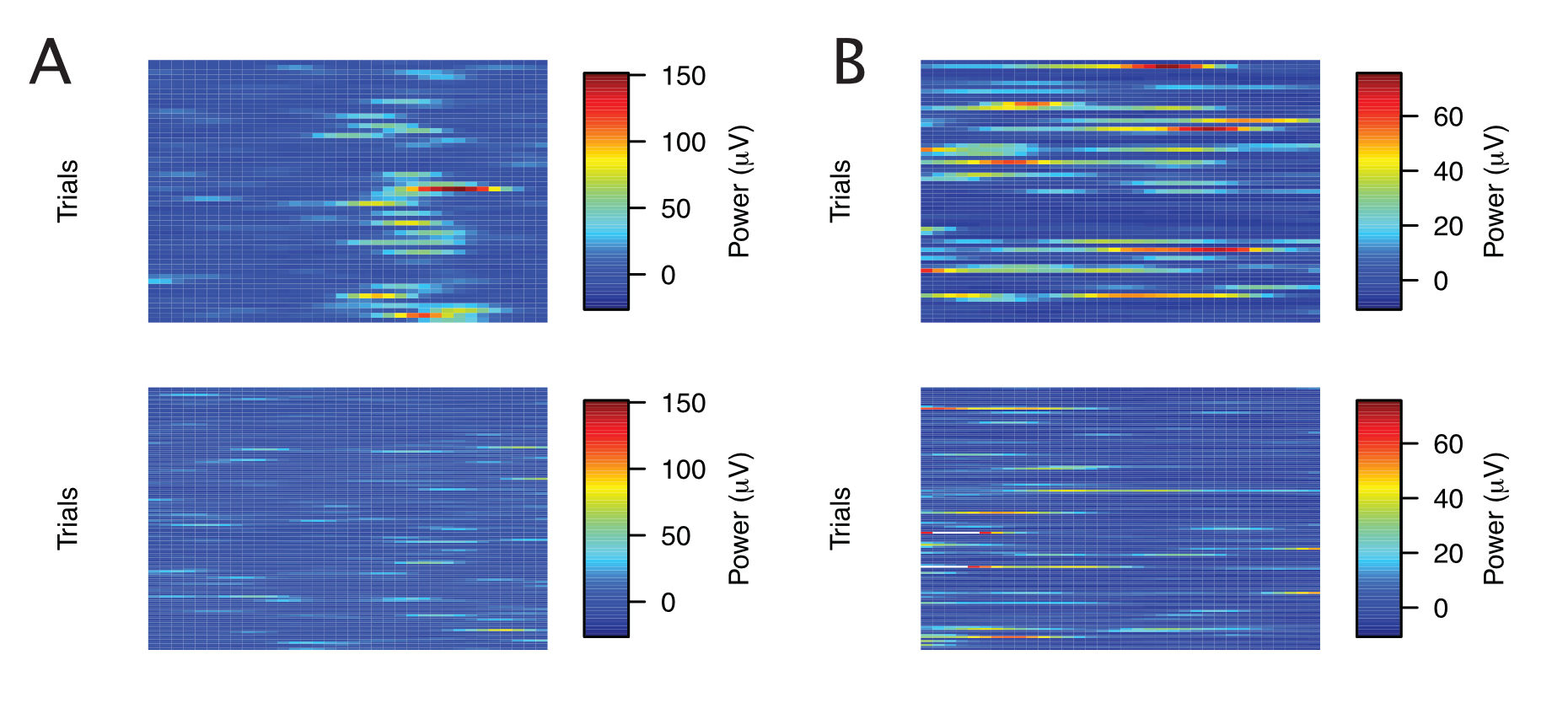

**Fig. S3 – Outcome encoding is associated with an increase in power.** (A) Distribution of T-stats for the association between gains and theta power. Positive values of the t-stat (to the right of the dashed red line) indicate higher power for gain than other trials, whereas negative values indicate the opposite. (B) As (A), but for loss encoding in beta.

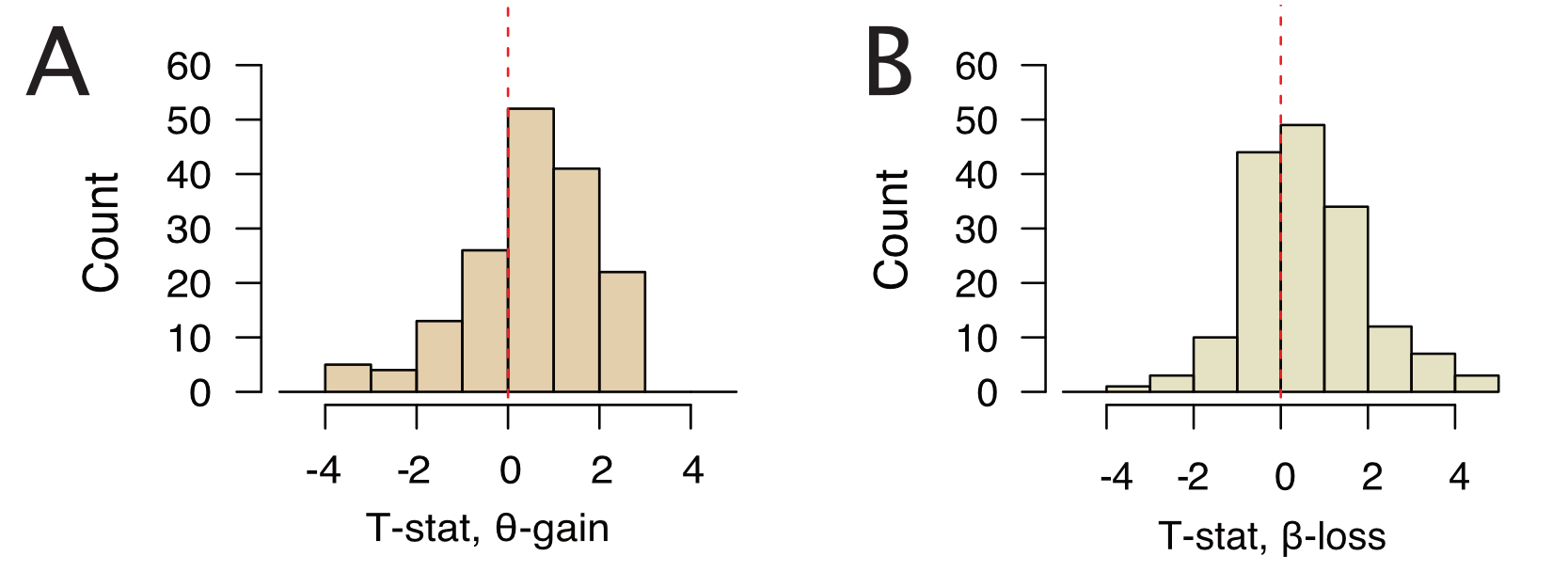

**Fig. S4 – Lack of anatomical segregation of theta-win and beta-loss encoding.** Electrode coordinates were cross-referenced with labeled anatomical atlases (JuBrain) to obtain their anatomical location according to Brodmann areas, verified by visual confirmation of electrode location based on surgical notes. We did not observe clear evidence for over or underrepresentation of beta or theta-encoding electrodes in Brodmann areas A11/A12/A13 (all p>0.05, Chi-square test). A10 and A14 were not tested due to the small proportion of electrodes in them.

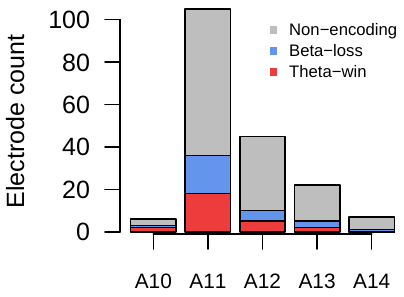

**Fig. S5 – Time courses of beta/theta encoding and coherence increases.** Plot shows the timecourse of Z-scored encoding (EV) and coherence, referenced to the outcome reveal time (t=0). Timecourses are shown separately for losses (blue) and gains (red). Both encoding and coherence are obtained from a single relevant frequency band: losses in beta band (maximum at 20Hz), and gains in theta band (max at 4Hz).

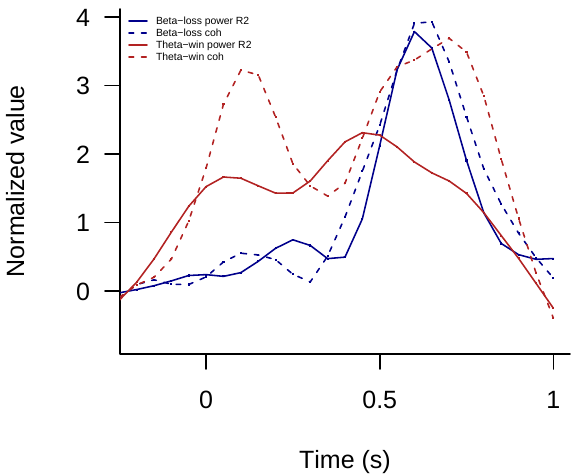
